# *In vivo* Morphological Dynamics of Single Laser-Axotomized Corneal Nerve Fibers in Sarm1-Null Adult Mice

**DOI:** 10.1101/2025.01.14.632447

**Authors:** Almudena Íñigo-Portugués, Fernando Aleixandre-Carrera, Salvador Sala, Fernando Borrás, Jose A. Gomez-Sanchez, Peter Arthur-Farraj, M. Carmen Acosta, Carlos Belmonte, Juana Gallar, Víctor Meseguer

## Abstract

Axonal regeneration represents a pivotal aspect of the adult peripheral nervous system (PNS). When an injury occurs, peripheral axons are able to regenerate and reestablish connections with their original targets. Although several precision techniques for severing axons in various non-mammalian model organisms have allowed to study the dynamics of regeneration after injury, similar axonal injury models have yet to be fully developed in mammals. In this study, we introduced a novel experimental method for laser-induced axotomy of single corneal sensory subbasal nerve fibers. This method enables the *in vivo* monitoring and quantification of axonal degeneration and regeneration morphological dynamics in adult mice. Results revealed that the degeneration of the distal stump of the subbasal fiber was delayed in mice lacking the protein SARM1, which promotes degenerative process after injury. Meanwhile, the proximal stump maintained the regenerative dynamics of the subbasal fiber, but regeneration of its nerve terminals was impaired. The present study introduces a valuable model for the *in vivo* study of the morphological plasticity dynamics of peripheral axons after injury in genetically tractable adult mammals.

## Introduction

Experimental models aimed at inducing controlled and reproducible injuries in mammals have uncovered key cellular and molecular aspects of the peripheral axon response to traumatic injury (Chen et al., 2007; Coleman & Freeman, 2010; Mahar & Cavalli, 2018; Wood et al., 2011). However, these models are constrained by their low regenerative capacity and offer limited information on the morphological dynamics of the axonal plasticity following injury (Griffin et al., 2010; Soares et al., 2014). While fixed preparations provide only limited insights into the morphological dynamics of the severed axon, *in vitro* paradigms can artificially influence the extent of morphological changes observed (Canty et al., 2013) and fail to fully capture the roles of surrounding cells in regulating axon degeneration (Mutschler et al., 2023) and regeneration (Arthur-Farraj et al., 2012). Consequently, live animal experimentation is essential to comprehensively elucidate the underlying mechanisms of post-injury axon plasticity (Villegas et al., 2012). Nevertheless, controlling the extent of the lesion and inflammation remains challenging and can significantly confound assessments of peripheral axonal degeneration and regeneration in surgery-induced injury paradigms (Canty et al., 2013).

Laser-induced local ablation of single axons has significantly overcome the aforementioned technical limitations in non-mammalian animal models. As such, the zebrafish (Villegas et al., 2012), the *C. elegans* (Bejjani & Hammarlund, 2012; Fatih Yanik et al., 2004) and the fruit fly *Drosophila melanogaster* (Fang et al., 2013; Soares et al., 2014) provide unique advantages such as optical transparency, genetic manipulation, the availability of laser-based methods to axotomize single axons and live imaging of their de-and regenerative processes. However, the lack of a similar injury paradigm in mammals hinders the extrapolation of results to humans from a more closely related evolutionary organism.

Therefore, we have established a pioneering model for inducing localized peripheral axonal injuries on individual nerve fibers. For this purpose, single subbasal fibers (SBFs) at the corneal epithelium were axotomized in living adult mice using a femtosecond laser. This enables *in vivo* monitoring of the severed fiber and subsequent quantification of the axonal degeneration and regeneration morphological dynamics (hereinafter referred to as morphodynamics). After axotomy, the distal stump of the subbasal fiber, along with their nerve terminals (SBF-NTs), experiences complete Wallerian degeneration (WD) within hours. In contrast, the proximal fiber stump undergoes rapid regeneration within days.

In recent years, the SARM1 protein (sterile-α and Toll/interleukin 1 receptor motif-containing protein 1) has emerged as a crucial intrinsic molecular mechanism in the WD of peripheral axons (Essuman et al., 2017; Gerdts et al., 2013, 2015; Osterloh et al., 2012). The SARM1 protein remains inactive in an intact peripheral neuron while it becomes activated upon axonal injury (Coleman & Höke, 2020; Gerdts et al., 2015; Gilley et al., 2015). Experiments involving SARM1-deficient mice have shown a significant delay in the degradation process of peripheral axons following an injury in mouse DRG sensory neurons (Geisler et al., 2019; Gerdts et al., 2015; Osterloh et al., 2012). Further, Sarm1 is required for regeneration of sensory and motor axons in injured sciatic nerves (Schmitd et al., 2024). Nevertheless, direct evidence for the influence of SARM1 on the morphological time dynamics of regeneration of severed peripheral axons is still unknown.

The present study demonstrates that individual corneal subbasal fibers lacking SARM1 undergo delayed degeneration after axotomy. The morphodynamics of the subbasal fiber regeneration is maintained, although the recovery of the nerve terminals is impaired.

## Results

### Selective laser ablation of individual subbasal fibers

To create precise lesions on peripheral axons, single fibers of the subbasal plexus at the corneal epithelium were reproducibly axotomized in anesthetized mice using a femtosecond laser coupled to a confocal microscope (Fig. 1A). We took advantage of the Trpm8^BAC^-EYFP mouse line, which allows visualization of individual nerve fibers by expressing the yellow fluorescent protein EYFP under the promoter of the gene for the cold-sensing ion channel TRPM8, in a reduced fraction, around 20 % of the total corneal innervation (Fig.1B) (Alcalde et al., 2018).

**Figure 1.**
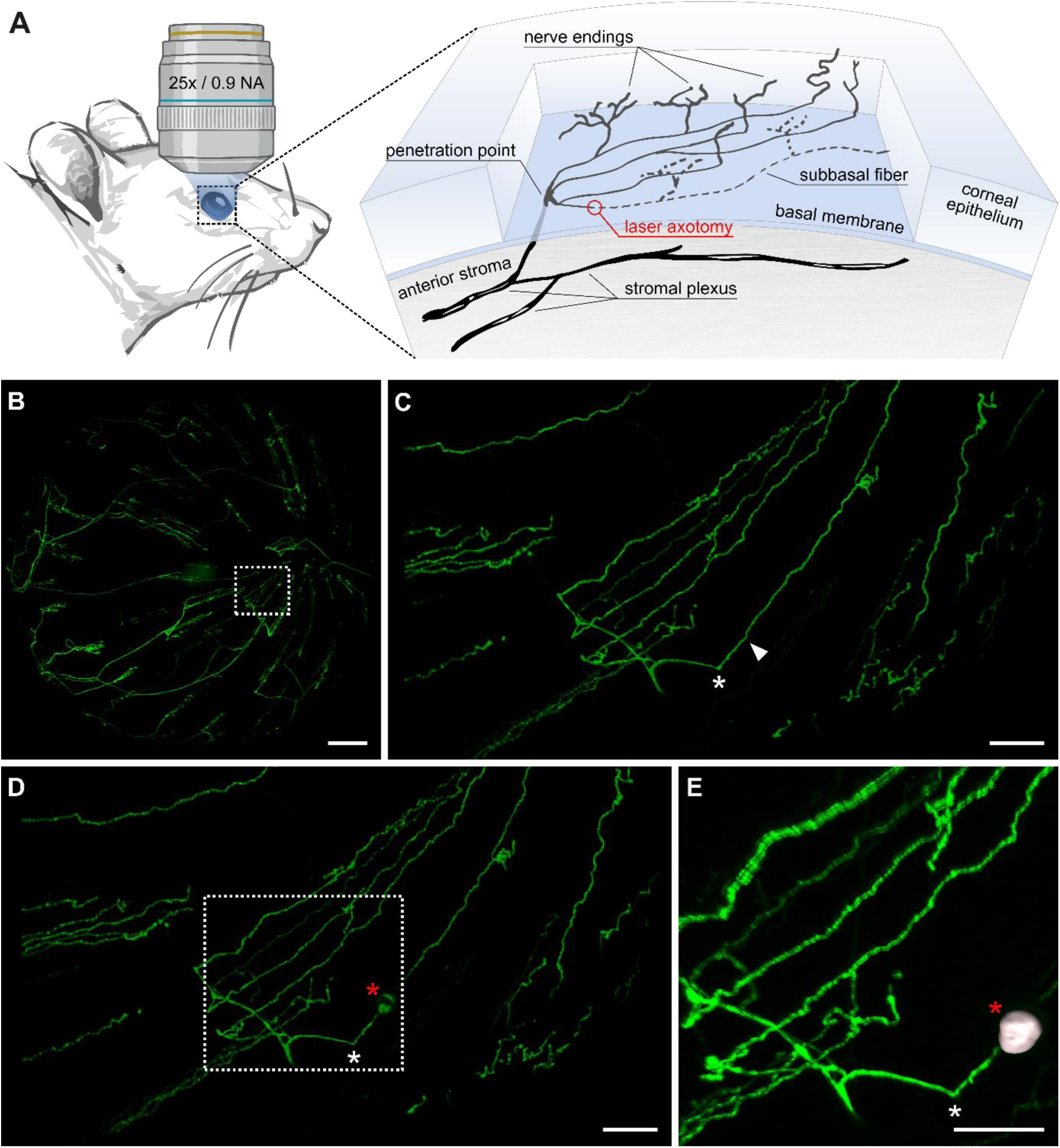
*In vivo* laser axotomy of a single subbasal fiber (SBF) in an adult mouse. **A.** Illustration of the laser-generated axotomy of a single SBF at the mouse corneal epithelium with the degenerating distal stump indicated by the dashed line. **B.** TRPM8^BAC^-EYFP corneal innervation in an intact cornea. The region delineated by the dotted box represents the epithelial leash where laser axotomy was performed and is magnified in (C). **C**. The epithelial leash was composed of five subbasal fibers. The penetration point is indicated with an asterisk, and the individual SBF selected for laser axotomy is indicated with an arrowhead. **D**. The SBF immediately after its laser axotomy. The red asterisk indicates the fluorescent mark. The dotted box highlights the area enlarged in (E). **E**. Three-dimensional reconstruction of the volume occupied by the fluorescent mark (in gray), superimposed on the original image. Scale bars: 300 μm in B and 50 μm in C-E.

After performing individualized laser ablation on the SBF (Fig 1B-E), a fluorescent mark was identified at the site of the lesion (Fig. 1D, E). This mark has previously been utilized to estimate the size of punctate lesions induced by a femtosecond laser in cortical neurons in the mouse (Canty et al., 2013). Then, we quantified the neural damage as a result of the laser ablation by measuring the volume of the fluorescent mark (Fig. 1E). The average volume was (5.16 ± 1.22) × 10^3^ µm^3^ (n = 9 fibers) which is similar to the volume of an average mammalian cell (Luby-Phelps, 2000). The observation revealed that SBFs separated by just a few micrometers from severed fibers maintained their original architecture. (Fig. 1D, E).

The data suggest that the lesions were highly precise at the target subbasal fiber.

### Assessment of the lesion damage at the corneal epithelium

To quantify the extent of the damaged tissue surrounding the microlesion, we fixed and immuno-stained the cornea with the cell nuclei dye Hoescht 33342 eight hours after the laser ablation (Fig.2). Despite the absence of apparent disappearance of cells surrounding the lesion, the nuclei of these cells exhibited a distinct elongated morphology around the fluorescent mark, differing from that observed in other cell nuclei located further away (Fig.2A). Furthermore, the cells near the SBF lesion showed an increased Hoechst 33342 emitted fluorescence intensity (Fig. 2A), which is indicative of cell damage (Crowley et al., 2016). Notably, the fluorescent mark is present exclusively in the epithelial layer of the lesion without affecting the adjacent corneal stroma (Fig. 2B). Next, we quantified the number of cell nuclei in proximity to the lesion revealing signs of morphological changes (Fig. 2C-E). As a result, we identified a total of 23.40 ± 2.27 cells (n=5 lesions) in the lesion’s surrounding area that exhibited higher fluorescence emission than the average of cells labeled with Hoescht 33342.

**Figure 2.**
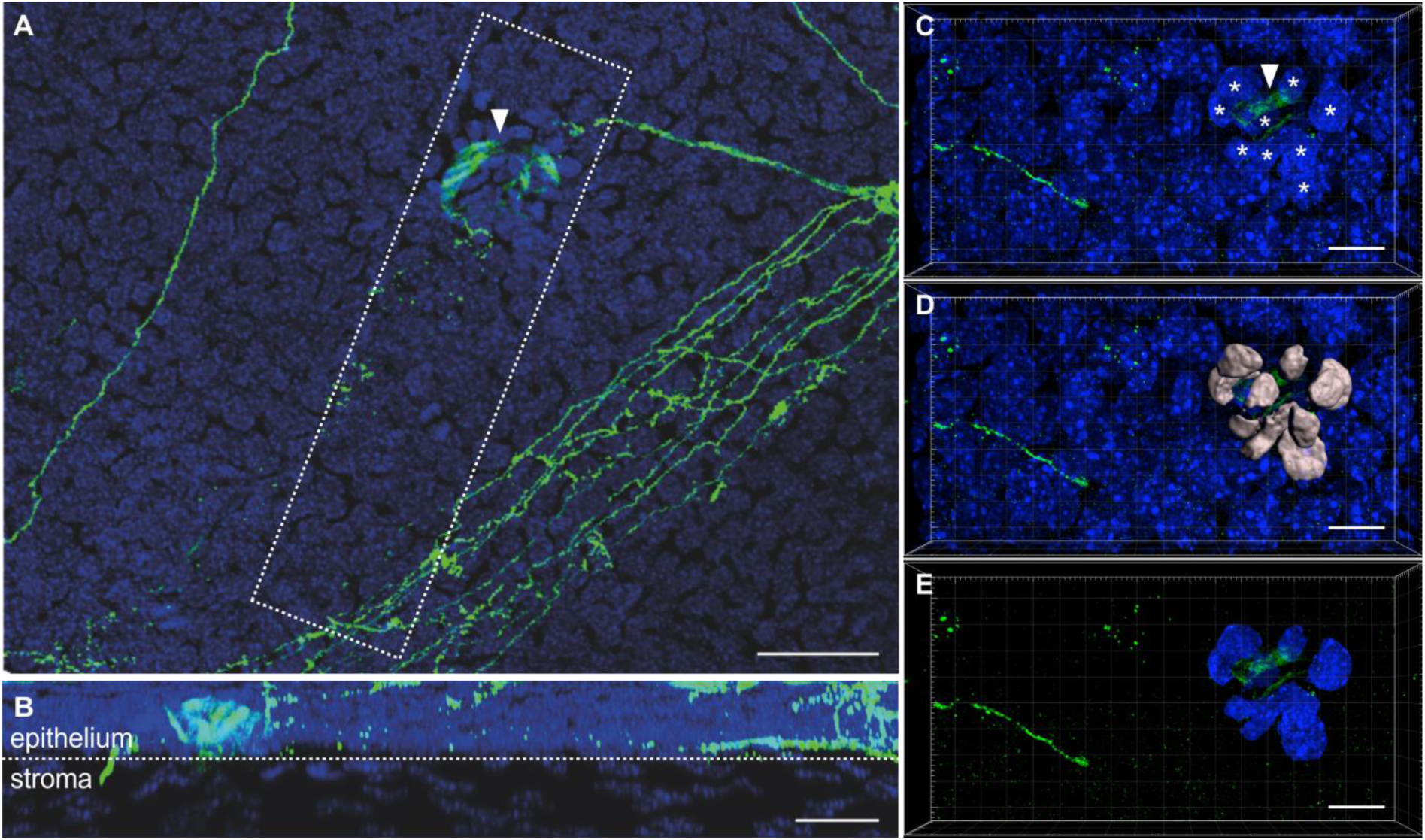
Potentially damaged cell nuclei surround the fluorescent mark at the corneal epithelium. **A.** Confocal image of the region where laser axotomy on a SBF was conducted. The cornea was fixed eight hours after the photoablation procedure. The nuclei of corneal cells were stained with Hoechst 33342. The arrow denotes the fluorescent mark. **B**. Orthogonal projection of the area indicated by a dotted rectangle in (A). The dotted line indicates the boundary between the epithelium and stroma. **C**. A three-dimensional representation of the region of the photoablation. The arrowhead indicates the fluorescent mark generated by the laser axotomy. Nuclei exhibiting increased fluorescence are marked with an asterisk. **D**. A three-dimensional reconstruction of damaged nuclei using Imaris 9.3 software (gray) overlaid on the original image. **E**. A modified image enables visualization of only the reconstructed nuclei. Scale bars represent 30µm in A and 10 µm in B-E.

These data indicate that the lesions had a markedly limited impact on the surrounding tissue.

### Rapid degeneration of the laser axotomized subbasal fiber and its nerve terminals in TRPM8^BAC^-EYFP mice

The laser axotomy (LA) caused the severed SBF to split into two segments. The distal stump - comprising the axotomized subbasal fiber and its nerve terminals (SBF-NTs) - extended from the fluorescence mark to the fiber’s peripheral end, while the proximal stump ranged from the fluorescence mark to the penetration point (Fig. 3A). Immediately after the laser axotomy, the intensity of EYFP fluorescence at the distal stump was not different from that before the lesion (Fig. 3A, F). Noteworthy, the fluorescence emitted by the EYFP protein in the distal stump of the severed control Trpm8^BAC^-EYFP/Sarm1^+/+^ fiber (n = 4 fibers) was absent 8 hours after the laser axotomy (Fig. 3F, 4A-D).

**Figure 3.**
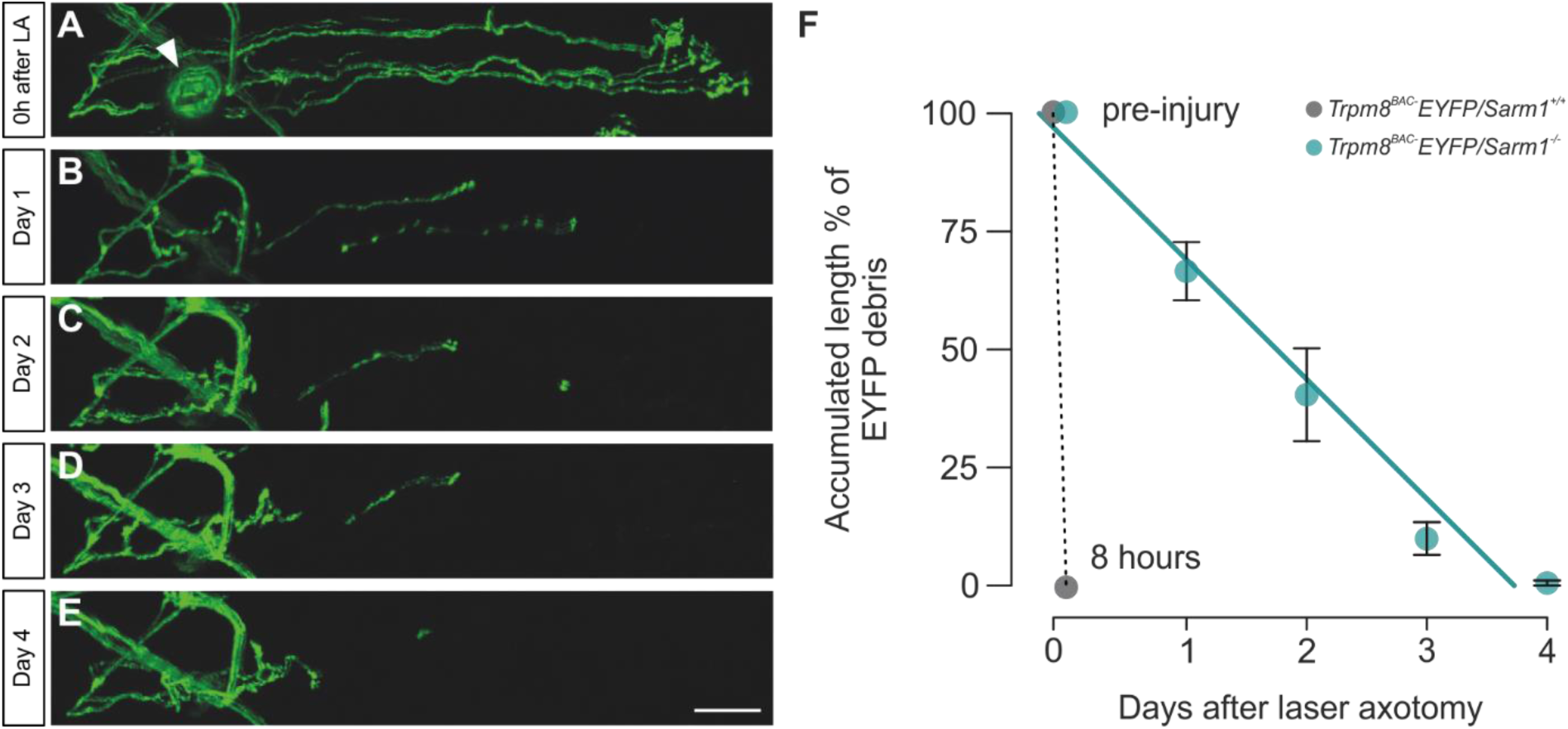
Dynamics of degeneration of the SBF and nerve terminals (NTs) following laser axotomy. **A.** *In vivo* confocal images of SBFs and their NTs immediately following laser ablation. The subsequent images in the series, **B** through **E**, show the situation on days 1, 2, 3, and 4, respectively. The arrow indicates the location of the laser ablation fluorescent mark. Scale bar in A-E: 50 µm. **F**. Decline in the proportion of accumulated EYFP residues over time. The circles represent the mean cumulative length for Trpm8^BAC^-EYFP/Sarm1^+/+^ (gray) and Trpm8^BAC^-EYFP/Sarm1^-/-^ fibers (green). The dotted line illustrates the rapid decay (within eight hours) of EYFP fluorescence in the distal stump of axotomized Trpm8^BAC^-EYFP/Sarm1^+/+^ fibers. The green line corresponds to the linear regression (R²=0.97, p<0.002) for the fluorescence decay of the Trpm8^BAC^-EYFP/Sarm1^-/-^ degenerating distal stumps.

These findings suggest the Wallerian degeneration process occurs in the severed SBF-NTs within the first hours after axotomy.

### Loss of Sarm1 delays degeneration of the subbasal fiber and its nerve terminals after laser axotomy

Recent studies show that the lack of the pro-degenerative protein SARM1 delays the degeneration of peripheral nerve fibers after their lesion in mice (Geisler et al., 2019; Gerdts et al., 2015). Here, we further investigated the involvement of SARM1 in the distal stump morphodynamics of degeneration at individual SBFs after their axotomy. Unlike control Trpm8^BAC^-EYFP/Sarm1^+/+^ fibers, the EYFP fluorescence remnants from the Trpm8^BAC^-EYFP/Sarm1^-/-^ distal stump were observed up to 4 days after the lesion (Fig. 3B-E, F). As such, the rate of disappearance of the accumulated length of the EYFP debris was 25.55% per day (n = 3 fibers; Fig. 3F).

Immunostaining of EYFP was performed on fixed corneas to determine whether the loss of that fluorescence is due to a reduction in EYFP fluorescence emission or the actual disappearance of the EYFP protein (Fig.4). In this manner, it was observed that in control Trpm8^BAC^-EYFP/Sarm1^+/+^ mice, there were no remnants of EYFP in the distal stump after 8 hours (Fig. 4C, D) while remnants of EYFP protein were observed in Trpm8^BAC^-EYFP/Sarm1^-/-^ mice (Fig. 4G, H).

**Figure 4.**
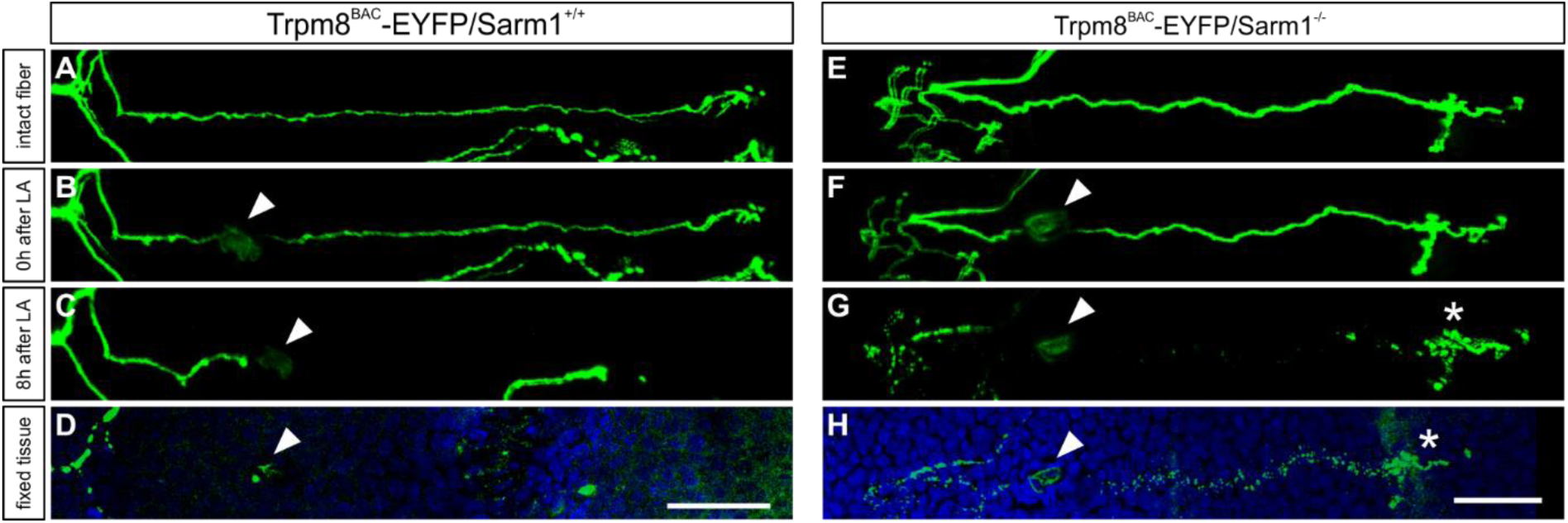
Persistence of the EYFP protein in the degenerating SBF and NTs following laser axotomy. *In vivo* confocal images of a control Trpm8^BAC^-EYFP/Sarm1^+/+^ (**A-C**) and a Trpm8^BAC^-EYFP/Sarm1^-/-^ fiber (**E-G**), before (A, E), immediately after (B, F), and 8 hours following laser axotomy (C, G). Afterwards, immunohistochemical analysis was performed in the Trpm8^BAC^-EYFP/Sarm1^+/+^ (**D**), and the Trpm8^BAC^-EYFP/Sarm1^-/-^ (**H**) subbasal fiber using anti-GFP (green) antibodies. Cell nuclei were stained with Hoechst 334442. The arrow heads indicate the location of the fluorescent mark generated by the laser photocoagulation. The asterisk denotes the remaining residues of EYFP in the degenerating distal stump of the fiber from the SARM1-lacking mouse. Scale bars: 50 μm.

These data indicate that the absence of the SARM1 protein dramatically delays the Wallerian degeneration process of the severed SBF-NTs after their laser axotomy.

### Regeneration morphodynamics of the axotomized subbasal fiber and its nerve terminals in TRPM8^BAC^-EYFP mice

One day after the laser axotomy, the proximal stump - comprising the regenerating SBF-NTs - exhibited a modest regrowth of 5.24 ± 3.69% (Fig.6A), as determined by the recovered cumulative length relative to its pre-injury value while partially recuperated up to 47.01 ± 12.24% on day 10 (Fig. 5E). The rate of regrowth was adjusted to an exponential growth curve (R² = 0.98) with an asymptote of 64.04%. This finding indicates that the SBF-NTs had reached maximum recovery by day 10, as predicted by the mathematical model. (Fig. 6A). The growth of the SBF and its NTs was then examined separately, in the horizontal plane (shown in green in Fig.5) which characterizes the subbasal fiber, and in the vertical plane (shown in white in Fig.5), corresponding to the nerve terminals (Marfurt et al., 2010). The analysis revealed a retraction (−5.00 ± 2.15%) of the remaining SBF (Fig. 6B) at day 1 post-injury. In comparison, the NTs length experienced a significant recovery (20.59 ± 5.72%) in the same period (Fig. 6C). On the third day following laser axotomy, the SBF length exhibited a modest enhancement (Fig. 5C), with a recovery of 3.30 ± 2.37% and a reach of 26.05 ± 6.44% by day 10 (Fig. 6B). However, the SBF length did not follow an asymptotic growth rate (R^2^ = 0.88) within the observed time frame (Fig. 6B). Remarkably, the NTs length recovered up to 77.62 ± 16.01% on day 3 and fully recovered the pre-injury length on day 10 (104.60 ± 23.80%), with an asymptote of rate regrowth of 108.40% (R^2^=0.98) (Fig. 6C).

**Figure 5.**
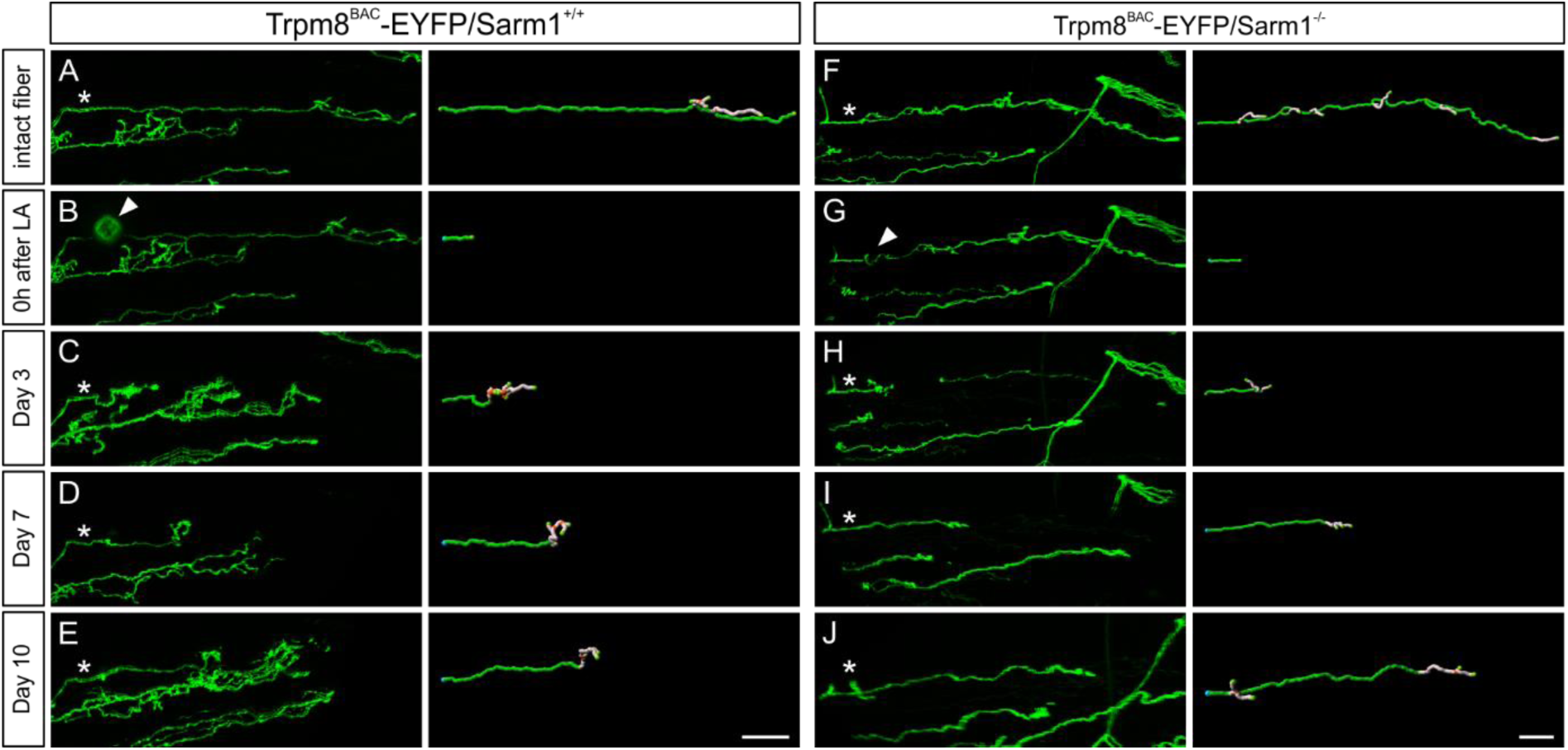
Dynamics of regeneration of the SBF and NTs following laser axotomy. *In vivo* confocal images of a Trpm8^BAC^-EYFP (**A-E**, left panels) and a Trpm8^BAC^-EYFP/Sarm1^-/-^ (**F-J**, left panels) subbasal fiber before their laser axotomy, immediately after (B, G), at day 3 (C, H) at day 7 (D, I), and at day 10 (E, J). The asterisks indicate the single fiber that was axotomized and monitored over time. A three-dimensional reconstruction of the severed corneal nerves of the Trpm8^BAC^-EYFP mouse (A-E, right panels) and the Trpm8^BAC^-EYFP/Sarm1^-/-^ mouse (F-J, right panels) shows the regenerating SBF in green and the NTs in gray. The fluorescent mark generated following laser axotomy is indicated by the arrowheads (B, G). The reconstruction of the distal stump in B and G is not shown to facilitate the tracking of regeneration. Scale bars: 50 µm.

**Figure 6.**
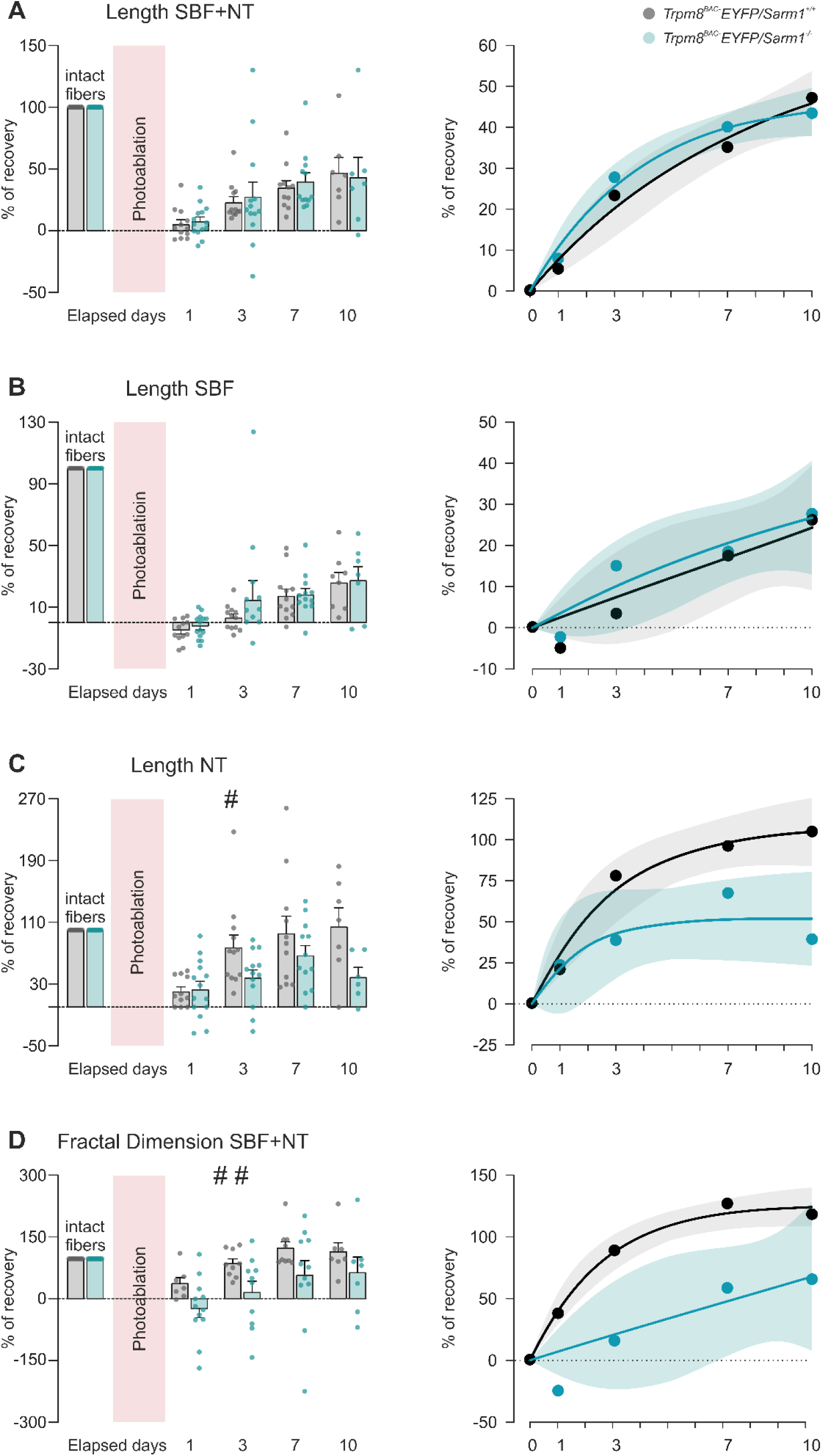
Comparison of the morphological regeneration dynamics of Trpm8^BAC^-EYFP and Trpm8^BAC^-EYFP/Sarm1^-/-^ SBFs and NTs following laser axotomy. Bar charts displaying the mean and standard error of the recovery (%) of severed fibers at the SBF+NT length (**A**, left), the cumulative length of their SBFs (**B**, left), the cumulative length of the NTs (**C**, left), and the fractal dimension of SBFs+NTs (**D**, left). The individual parameter values for each Trpm8^BAC^-EYFP (grey dots) and Trpm8^BAC^-EYFP/Sarm1^- /-^ (green dots) axotomized fiber are represented as points on the bars. The orange bar indicates the moment of photocoagulation. The horizontal dotted line represents the baseline. A mixed-effects analysis was conducted, followed by Sídák’s test for multiple comparisons with respect to the values observed prior to photocoagulation. # (p<0.05) and ## (p<0.01) denotes statistical significance due to the global observed differences according to genotype and not referring to a particular time point. The right panels illustrate the dynamics of recovery of the SBF+NTs length (A), SBF length (B), NTs length (C), and the fractal dimension of SBF + NTs (D), expressed as a percentage of the pre-injury values. These values were obtained by fitting the mean values at different time points to an exponential growth curve constrained to cross the origin of coordinates. Shaded areas represent confidence intervals of the fitted curves.

The recovery of the morphological complexity of the regenerating SBF-NTs was quantified by measuring its fractal dimension (FD) (Chen et al., 2018; Petropoulos et al., 2020). A significant recovery of the SBF-NT fractal dimension (37.60 ± 14.89%) was observed as soon as one day after the axotomy. This recovery was not significantly different from its pre-lesion value at day 3 post-injury (88.42 ± 11.53%) (Fig. 6D). Throughout the 10-day period, the rate of regrowth of the SBF-NT fractal dimension reached an asymptote at 126.60% (R^2^=0.99) (Fig.6D).

These results demonstrate a regeneration process in the severed subbasal fiber and its nerve terminals, evident on the initial day following the injury. A discernible disparity in the recovery of the nerve terminals and the subbasal fiber is evident post-injury, with a rapid and complete regeneration of the nerve terminals’ cumulative length and the complexity of the subbasal fiber and their nerve terminals. In contrast, the regrowth of the subbasal fiber alone is slower and incomplete.

### Impaired regeneration of nerve terminals lacking SARM1

Following laser axotomy, the proximal stump of Trpm8^BAC^-EYFP/Sarm1^-/-^ subbasal fibers showed a partial recovery, exhibiting morphodynamics of regeneration that were comparable to Trpm8^BAC^-EYFP fibers. This is evidenced when comparing the SBF-NTs length (R^2^= 0.99; asymptote of 47.23%) (Fig. 5G-J; 6A) and SBF length (R^2^= 0.90; asymptote= 51.87%) (Fig. 5G-J; 6B). In contrast to the complete regeneration observed in severed Trpm8^BAC^-EYFP fibers (see Fig. 6C), the Trpm8^BAC^-EYFP/Sarm1^-/-^ proximal stumps displayed only partial recovery in the NTs length (R^2^= 0.81; asymptote of 52.03%) and the fractal dimension of the SBF-NTs (R^2^= 0.80) (Fig. 6D).

This set of data suggests that while the absence of SARM1 does not impact the morphodynamics of the regenerating subbasal fiber cumulative length, it does impede the morphodynamics of their regenerating nerve terminals, also having an impact on the complexity of the subbasal fiber and its nerve terminals as a whole.

## Discussion

This study quantifies, for the first time in mammals, the morphological dynamics of local degeneration and regeneration of peripheral axons *in vivo* after laser axotomy of individual corneal subbasal fibers in mice. Most studies on peripheral nerve regeneration have focused on lesions resulting in partial or complete loss of a large number of peripheral axons in rodents (Griffin et al., 2010; Song et al., 2019; Wood et al., 2011). Similarly to the zebrafish pLL nerve (Villegas et al., 2012) and drosophila nerve wing (Soares et al., 2014) laser axotomy paradigms, here we axotomized thin individual corneal subbasal fibers, each containing a few axons as shown by microscope electron studies in mice and humans (∼ 10 axons) (Müller et al., 2003; Whitear, 1960).

As a result of the laser axotomy, a small fluorescent mark was generated at the severed single subbasal fiber, reflecting the small size of the neural lesion. Notably, no changes were observed in the adjacent subbasal fibers to the severed fiber immediately following laser axotomy. The data presented here are consistent with a previous report (Allegra Mascaro et al., 2013), in which single axons in mouse cerebellar climbing fibers were ablated using laser axotomy, thus avoiding collateral damage to adjacent dendrites.

The subbasal fibers are located between the basal and suprabasal cells of the corneal epithelium. No intercellular spaces corresponding to those presumably left by a disrupted tissue were observed after laser axotomy. However, Hoechst staining, a fluorescent marker commonly used to visualize cellular DNA damage (Crowley et al., 2016), reveals evidence of damage in several cells surrounding the fluorescent marker. Laser axotomy of the subbasal fiber was always performed near the penetration point, just a few micrometers above the basal lamina, with no evidence of tissue damage in the stromal region closest to the lesion. These data confirm the finding of a highly localized lesion in the subbasal fiber, with minimal involvement of the surrounding epithelial tissue.

The rapid dynamics of postlesional plasticity of the severed subbasal fiber, on the time scale of days, suggests the existence of a permissive nerve regenerative environment. The absence of myelin and associated proteins in the subbasal fiber (Frutos-Rincón et al., 2022) may account for this, as they have been proposed to have an inhibitory effect on post-injury axonal regrowth (Filbin, 2003; Villegas et al., 2012). Furthermore, the rapid turnover of corneal epithelial cells (Harris & Purves, 1989) may contribute to the permissive environment by providing the proximal regenerating stump with growth alternatives to the spaces previously occupied by the degenerating distal portion of the corneal nerve (Buck, 1985; Harris & Purves, 1989; Sagga et al., 2018).

Interestingly, we found that fibers do not recover their morphological structure homogeneously. Nerve terminals recover their full initial cumulative length one week after axotomy, whereas the subbasal fiber recovers poorly within the same period. Moreover, our data show full recovery of the complexity of the regenerating fiber long before it reaches its pre-lesion size. Since TRPM8-expressing nerve terminals sense and encode the information about temperature decreases at the ocular surface, the early recovery of both nerve terminals’ cumulative length and the regenerating fiber complexity may contribute to their functional restoration. Consistent with this, Bech et al. (2018) recorded nerve terminal impulses evoked by cold from the regenerating corneal nerves at the injury site as early as 7 days after photorefractive keratectomy.

Schwann cells play a crucial role in post-injury peripheral nerve plasticity (Arthur-Farraj et al., 2012; Jessen & Mirsky, 2016). It is worth noting that while stromal nerve bundles are wrapped by non-myelinating corneal Schwann cells, subbasal fibers lack Schwann cells (Bargagna-Mohan et al., 2021; Müller et al., 1996, 2003) and are instead wrapped by corneal epithelial basal cells. Corneal epithelial cells have been reported to act as surrogate Schwann cells in the corneal epithelium since they express similar molecular markers (Stepp et al., 2017). However, further functional evidence is necessary to elucidate the contribution of corneal epithelial cells to axonal post-lesion morphological plasticity.

The SARM1 protein has been postulated as an intrinsic molecular mechanism essential in activating the Wallerian degeneration process that follows peripheral nerve injury (Osterloh et al., 2012). Injury of a peripheral axon activates the pro-degenerative SARM1 protein, promoting the Wallerian degeneration process in the distal portion of the severed axon (Gerdts et al., 2013, 2015; Osterloh et al., 2012). Here, we show that in control mice, the distal portion of the TRPM8 corneal subbasal fibers degenerates completely in hours after its photoablation, consistently with the rapid Wallerian degeneration observed in single injured axons in non-mammalian paradigms (Martin et al., 2010). In contrast, in mice lacking the SARM1 protein, a complete degeneration of the distal part is observed in the days following the lesion. These results are consistent with the delayed Wallerian degeneration reported in axotomized axons *in vitro* from mouse Sarm 1^-/-^ dorsal root ganglia sensory neurons and mouse Sarm1^+/−^ sciatic nerves lesioned *in vivo* (Osterloh et al., 2012).

The present study demonstrates that the absence of SARM1 does not affect the morphological dynamics of regeneration of the severed subbasal fiber while impairing their nerve terminal regeneration. The persistence of axon debris in the Sarm1^-/-^ mouse may contribute to the repeal of regenerating peripheral axons (Bisby & Chen, 1990; Brown et al., 1992; Martin et al., 2010). Similarly, a recent preprint demonstrates that Sarm1^-/-^ distal nerve tissue of the crushed mouse sciatic nerve exhibits a growth-inhibitory environment, impairing regeneration (Schmitd et al., 2024). Remarkably, the human SARM1, when expressed in a C. elegans model through transgenesis, exerts a regulatory function in the process of axonal regeneration in laser axotomized GABA motor neurons cell-autonomously (Czech et al., 2023). This finding indicates a possible role for SARM1 as an intrinsic mechanism of peripheral axonal regeneration.

Together, these results support the mouse individual subbasal fiber laser axotomy as a powerful model for live studies of axon degeneration and regeneration. This allows the *in vivo* study of the molecular and cellular mechanisms mediating post-injury morphological plasticity in adult mammals.

## Material and Methods

### Experimental animals

Experimental procedures were carried out on both male and female mice aged between 3 month and 6 months, following the institutional animal care guidelines outlined in the Spanish Royal Decree 53/2013 and the European Union Directive 2010/63/EU on the protection of animals used for experimental purposes. All the procedures were approved by the Ethics and Integrity in Research Committee of the Miguel Hernández University of Elche and authorized by the Conselleria of Agriculture, Rural Development, Climatic Emergency and Ecological Transition of the Generalitat Valenciana. The animals were bred and housed in the animal facility of the Miguel Hernández University of Elche in temperature-controlled rooms (21°C), 12-hour light-dark, and access to food and water ad libitum. For the development of the individual subbasal fiber laser axotomy and the quantification of their regenerative morphological dynamics, the Trpm8^BAC^-EYFP mouse line was used. This genetically modified mouse line expresses the EYFP protein under the control of the transcriptional regulatory sequences of the Trpm8 gene. Transgenesis was performed by modifying of a bacterial artificial chromosome containing the complete mouse Trpm8 locus and cDNA encoding the enhanced yellow fluorescent protein (EYFP) was inserted into it (Morenilla-Palao et al., 2014). The genetic modification permits the coexpression of TRPM8 and EYFP in primary sensory neurons, such that the excitation of the protein with light at a wavelength of 512 nm produces a fluorescence emission in those neurons that express TRPM8 natively. For experiments with corneal subbasal fibers lacking the SARM1 protein, a mouse line with a knockout genetic modification for the gene Sarm1 (Kim et al., 2007) was crossed with the Trpm8^BAC^-EYFP line to obtain Trpm8^BAC^-EYFP/Sarm1^-/-^ mice and their Trpm8^BAC^-EYFP/Sarm1^+/+^ littermates as control possessing a wild-type genotype for Sarm1.

### *In vivo* imaging of corneal innervation

Trpm8^BAC^-EYFP and Trpm8^BAC^-EYFP/Sarm1^-/-^ transgenic mice were anesthetized by intraperitoneal injection of xylazine (Xilagesic 16mg/kg). They were then head restrained in a mouse-specific mobile holder (SGM-4, Narishige International USA, Amityville, NY, USA) using bars placed over the left and right temporal bones. For the maintenance of a stable and prolonged degree of anesthesia in the mouse, volatile anesthesia was provided using an isoflurane vaporizer (FUJIFILM VisualSonics, Toronto, ON, Canada) with a built-in mask that allowed continuous inhalation of isoflurane (1-2%) during the procedure. The head holder was placed on an *in vivo* stage for intravital imaging experiments (Luigs&Neuman, Ratingen, Germany), with an XY motorization designed for the DM6000 CFS microscope (Leica Microsystems, Wetzlar, Germany). Subsequently, by manually adjusting the position of the movable mouse head holder, the ocular surface of the right eye was placed under the objective, allowing visualization of the fluorescent nerve fibers once they were excited with the corresponding laser. The left eye was continuously moistened by applying artificial tears (Optiben, Laboratorios Cinfa, Navarra, Spain) on the ocular surface to prevent evaporation of the ocular film and thus protect the corneal epithelium. A series of images were then acquired with a low magnification dry objective (5x) to determine the state of the corneal innervation, after which images of the fibers under study were acquired using a water immersion objective at higher magnification (25x). In this case, a drop of the artificial tear was applied between the corneal surface and the objective to allow visualization of individual subbasal fibers. Once the image acquisition was completed, the mouse head was released from the holder, injected subcutaneously with 1 ml of glucose solution and finally placed on top of a thermal blanket regulated at 37°C until recovery from anesthesia.

*In vivo* confocal imaging was performed using a LEICA SP5-II laser scanning microscope. A 5x dry objective (HCX PLFLUOTAR CS 5x/0.15) and a water immersion objective (HCX IRA APO L 25x/0.95 W). The EYFP protein was excited using an argon ion laser with light at a wavelength of 513 nm and the detection was carried out using a spectral photomultiplier between 525 and 600 nm. A series of z-stacked images of the fluorescent fibers was acquired using LAS AF software version 2.7.3.9723 (Leica Microsystems). The images were taken at a resolution of 1024×1024 pixels and with a depth of 8 bits. The scanning speed was 1200 Hz for all acquisitions. The size of each pixel was 2.90 x 2.90 μm for images acquired at 5x and 0.58 x 0.58 μm for those acquired at 25x. A z-step plane jump (z-step) of 9.68 μm for images acquired at 5x and a plane jump of 0.49 μm was set for the images acquired at 25x. All images were saved in Leica Image Format (.lif) for later analysis.

### *In vivo* laser axotomy of individual corneal subbasal fibers

Laser axotomy was performed on individual subbasal fibers using a femtosecond laser MaiTai HP DeepSee infrared laser (Spectra-Physics, Mountains View, CA) at a wavelength of 800nm, optically connected to the LEICA SP5-II laser scanning microscope. The infrared (IR) laser emits pulses at 100 femtoseconds interval, concentrating the photons at the focal (z) plane. The region of interest to be photo-coagulated was then selected, and a zoom-in was applied at a location close to the penetration point of the subbasal fiber. Then, an upper and a lower boundary was defined in the Z-plane to delimit the exact point where photocoagulation was to be performed, and a distance of 1 μm was established between the different optical sections (z-step) for image acquisition. The scanning speed was 1200 Hz and a double scan per line was configured (line average=2). After firing with the IR laser, a visual check with the microscope was performed to verify that the selected subbasal fiber had been photo-coagulated. This was possible because, after axotomy, a fluorescent mark was generated at the site where the laser had struck.

As previously described (Canty et al., 2013), the Imaris 9.3 surface tracing tool (Oxford Instruments) was used to reconstruct the fluorescence mark generated by the femtosecond laser and evaluate the volume of injured tissue after the laser axotomy (n=9) in the Trpm8^BAC^-EYFP mice.

Severed Trpm8^BAC^-EYFP and Trpm8^BAC^-EYFP/Sarm1^-/-^ subbasal fibers were monitored by image acquisition at days 0, 1, 3, 7 and 10 post-injury. During this period, the animals were individually housed. Those animals that experienced a loss of fluorescence in any region or in the totality of the corneal innervation, at any time point during the follow-up were sacrificed and their data excluded from the morphometric analysis.

### Morphometrics of single axotomized subbasal fibers and their nerve terminals

The cumulative length of the fibers was quantified from the three-dimensional reconstruction performed with Imaris 9.3 filament tracing tool (Oxford Instruments). The complexity of the fibers was estimated based on their fractal dimension value. To this end, the box-counting method (Barnsley, 1993; Falconer, 1997, 2003) was employed in a custom script developed in Python. A grid of boxes of varying sizes was superimposed over the subbasal fibers and their nerve terminals, and the number of boxes containing a portion of the object was counted. The FD was then calculated from the relationship between the number of boxes (N) and the size of the boxes (ε), where FD was the slope of the regression line obtained by plotting Log (N(ε)) versus Log (1/ε) at each iteration. The degree of regeneration of the injured corneal nerve fiber was quantified as a percentage of recovery relative to the preinjury value of cumulative length or fractal dimension, using the following equation:

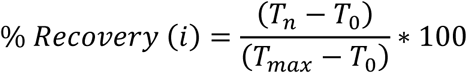

In this equation, *T_n_* represents the value of the cumulative length or the fractal dimension (*i*) on day *n* of the study. *T*_0_ denotes the value of (*i*) immediately following axotomy, while *T_max_* indicates the maximum value attained by (*i*) throughout the follow-up period.

### Immunohistochemistry

Trpm8^BAC^-EYFP and Trpm8^BAC^-EYFP/Sarm1^-/-^ mice used in the different experiments were sacrificed by intraperitoneal injection of sodium pentobarbital (Dolethal, Vetoquinol, Lure, France). Then, cervical dislocation was performed, the eyes were carefully enucleated and a puncture was made at the sclerocorneal limbus of the eyes to allow the fixation solution to penetrate the interior of the eye and the fixation to be homogeneous. The eyes were subsequently fixed for 2 hours in a solution of methanol and dimethyl sulfoxide (4:1) under agitation and at room temperature. Next, the cornea was dissected and four cuts were made to form a 4-petal clover to flatten the cornea. Immediately after, the corneas were post-fixed for 5 minutes in absolute methanol at -20°C and rehydrated in a decreasing battery of methanol with PBS. The corneas were washed 3 times in PBS- X at 0.1% for 30 minutes (10 minutes per wash) and incubated for 2 hours in blocking solution composed of 1% bovine serum albumin (Vector Laboratories) and 10% goat serum (Vector Laboratories), all diluted in PBS-X.

Subsequently, the corneas were incubated with chicken anti-GFP primary antibody (1:500; Abcam, Cambridge, MA, USA) in blocking solution for 2 days at 4°C. Then corneas were washed 5 times in PBS (10 minutes per wash), and the incubated with Alexa Fluor 488 goat anti-chicken IgG (ThermoFisher Scientific, Waltham, MA, USA) diluted in PBS (1:500) for 2 hours. Subsequently, 5 washes of 10 minutes each in PBS were performed, and then, the corneas were incubated with the DNA fluorescent dye Hoechst 33342 (Thermo Fisher Scientific) diluted in PBS (1:1000) for 10 min. This was followed by 3 washes of 10 minutes, and finally, the corneas were mounted on a slide using as mounting medium Fluoromount-G (ThermoFisher Scientific). Negative controls were made for all the used secondary antibodies by repeating the same immunostaining protocol, but without incubating the primary antibodies.

### Imaging of fixed corneal samples

All immunofluorescence images of nerve fibers in fixed corneas were acquired with a Zeiss LSM880 inverted confocal microscope (Carl Zeiss AG, Oberkochen, Germany). For this purpose, an objective with 25x oil immersion objective lens (Apochromat 25x/0.8 Imm DIC) and a 63x oil-immersion lens objective (Plan-Apochromat 63x/1.4 Oil DIC) were used. Hoechst 33342 and anti-GFP were excited with light at a wavelength of 405 nm and 488 respectively. The excitation and detection of the signal produced by the fluorochromes were performed using spectral photomultipliers and in a sequential way to avoid overlapping of the emission spectra between the different fluorophores. Fluorescence images were acquired at a resolution of 1024×1024 pixels and 8-bit depth using Zen Black version 2.3 software (Carl Zeiss AG). The scanning radius was 2.05 μs/pixel when using the 25x objective and 8.19 μs/pixel when the 63x was used. The size of each pixel is 0.33×0.33 μm in all the images acquired. The plane hopping (z-step) was defined according to the Nyquist-Shannon sampling theorem as 1.39 μm for images acquired at 25x and 1.15 μm for those acquired at 63x. All images were saved in Carl Zeiss Image (.czi) data format for further analysis.

### Statistics and analysis

The data were statistically analyzed using GraphPad Prism version 9.0.1 (GraphPad Software, La Jolla, CA, USA) and IBM SPSS Statistics for Windows, version 22 (IBM Corporation, Armonk, NY, USA). All results were expressed as the mean value ± standard error of the mean.

The statistical differences between variables were assessed using mixed-effects analysis followed by Sídák’s correction for multiple comparisons. Data was also fitted to an exponential growth curve to evaluate the time course evolution of the studied variables, according to the equation:

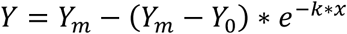

where *Y_m_* represents the maximum asymptotic value, *Y*_0_ the starting value and *k* the rate of growth constant. Growth curves were constrained by setting the parameters *Y*_0_ = 0 and *k* > 0, which we considered appropriate according to the experimental design. P-values < 0.05 were considered statistically significant. All the statistical tests used in each case are indicated in the text of their corresponding Results section and in the figure captions.

## Acknowledgements

This work was supported by the Spanish Ministry of Science and Innovation-Agencia Estatal de Investigación (MCIN/AEI/ 10.13039/501100011033) with grants PID2021-124460OB-I00 (V.M.) and PID2020-115934RB-I00 (J.G. & M.C.A.). Grant CIPROM/2021/48 from the Generalitat Valenciana, Spain, is also acknowledged. PA-F (206634/Z/17/Z) was funded by the Wellcome Trust (UK). For the purpose of Open Access, the author has applied a CC BY public copyright license to any Author Accepted Manuscript version arising from this submission. JAGS research was supported by a Miguel Servet Fellowship from the Spanish Health Institute Carlos III (CP22/00078).

